# Fine-scale computations for adaptive processing in the human brain

**DOI:** 10.1101/2020.02.14.947895

**Authors:** E Zamboni, VG Kemper, NR Goncalves, K Jia, VM Karlaftis, SJ Bell, JJ Giorgio, R Rideaux, R Goebel, Z Kourtzi

## Abstract

Adapting to the environment statistics by reducing brain responses to repetitive sensory information is key for efficient information processing. Yet, the fine-scale computations that support this adaptive processing in the human brain remain largely unknown. Here, we capitalize on the sub-millimetre resolution afforded by ultra-high field imaging to examine BOLD-fMRI signals across cortical depth and discern competing hypotheses about the brain mechanisms (feedforward vs. feedback) that mediate adaptive visual processing. We demonstrate suppressive recurrent processing within visual cortex, as indicated by stronger BOLD decrease in superficial than middle and deeper layers for gratings that were repeatedly presented at the same orientation. Further, we show dissociable connectivity mechanisms for adaptive processing: enhanced feedforward connectivity within visual cortex, while feedback occipito-parietal connectivity, reflecting top-down influences on visual processing. Our findings provide evidence for a circuit of local recurrent and feedback interactions that mediate rapid brain plasticity for adaptive information processing.

## Introduction

Interacting in cluttered and complex environments, we are bombarded with plethora of sensory information from diverse sources. The brain is known to address this challenge by reducing its responses to repeatedly or continuously presented sensory inputs. This type of sensory adaptation is a rapid form of plasticity that is critical for efficient processing and has been shown to involve changes in perceptual sensitivity (for review: Clifford, 2002) and neural selectivity (for review: Kohn, 2007). Numerous neurophysiological studies (for review: Kohn, 2007) have shown sensory adaptation to be associated with reduction in neuronal responses that are specific to the features of the adaptor. Functional brain imaging studies in humans have shown fMRI adaptation for low-level visual features (e.g., contrast, orientation, motion; for review Larsson, Solomon, & Kohn, 2016) as indicated by decreased BOLD responses in visual cortex due to stimulus repetition. Similar BOLD decreases have been reported in higher visual areas for repeated presentation of more complex visual stimuli (e.g. faces, objects), an effect known as repetition suppression (Grill-Spector, Henson, & Martin, 2006; Krekelberg, Boynton, & van Wezel, 2006).

Despite the plethora of studies investigating the neural signatures of adaptation, the fine-scale human brain computations that underlie adaptive processing remain debated. In particular, neurophysiological studies focussing on primary visual cortex provide evidence of rapid adaptation at early stages of sensory processing (Gutnisky & Dragoi, 2008; Whitmire & Stanley, 2016; Xiang & Brown, 1998). In contrast, fMRI studies have suggested top-down influences on sensory processing of repeated stimuli via feedback mechanisms (e.g., Ewbank et al., 2011; Summerfield, Trittschuh, Monti, Mesulam, & Egner, 2008). Yet, the circuit mechanisms that mediate adaptive processing in the human brain remain largely unknown, as fMRI at standard resolution does not allow us to dissociate feedforward from feedback signals.

Here, we capitalize on recent advances in brain imaging technology, to discern the contribution of feedforward vs. feedback mechanisms to adaptive processing. Ultra-high field (UHF) imaging affords the sub-millimetre resolution necessary to examine fMRI signals across cortical layers in a non-invasive manner, providing a unique approach to interrogate human brain circuits at a finer scale (for review: Lawrence, Formisano, Muckli, & de Lange, 2019) than that possible by standard fMRI techniques (for review: Goense, Bohraus, & Logothetis, 2016). UHF laminar imaging allows us to test the finer functional connectivity across cortical layers based on known anatomical laminar circuits. In particular, sensory inputs are known to enter the cortex from the thalamus at the level of the middle layer (layer 4), while output information is fed forward from superficial layers (layer 2/3), and feedback information is exchanged primarily between deeper layers (layer 5/6) as well as superficial layers (for review: Self, van Kerkoerle, Goebel, & Roelfsema, 2019).

Here, we combine UHF laminar imaging with an orientation adaptation paradigm (i.e. observers are presented with gratings at the same or different orientations) to test whether orientation-specific adaptation alters input processing in middle V1 layers, or feedback processing in superficial and deeper V1 layers (Figure 1A). We demonstrate that adaptation alters orientation-specific signals across layers within primary visual cortex with stronger fMRI-adaptation (i.e. BOLD decrease for repeated stimuli) in superficial layers. These signals are read-out by higher visual areas, as indicated by enhanced feedforward connectivity between V1 superficial and V4 middle layers. Further, we test the role of the posterior parietal cortex in adaptive processing, as it is known to be involved in stimulus expectation and novelty detection (de Lange, Heilbron, & Kok, 2018; Garrido, Kilner, Stephan, & Friston, 2009; Li, Gratton, Yao, & Knight, 2010; Summerfield & De Lange, 2014). Our results show enhanced feedback connectivity from posterior parietal cortex to V1 deeper layers, suggesting top-down influences on visual processing via feedback mechanisms. Our findings provide evidence for a circuit of local recurrent processing across layers within visual cortex and occipito-parietal feedback interactions that mediate adaptive processing in the human brain.

**Figure 1.**
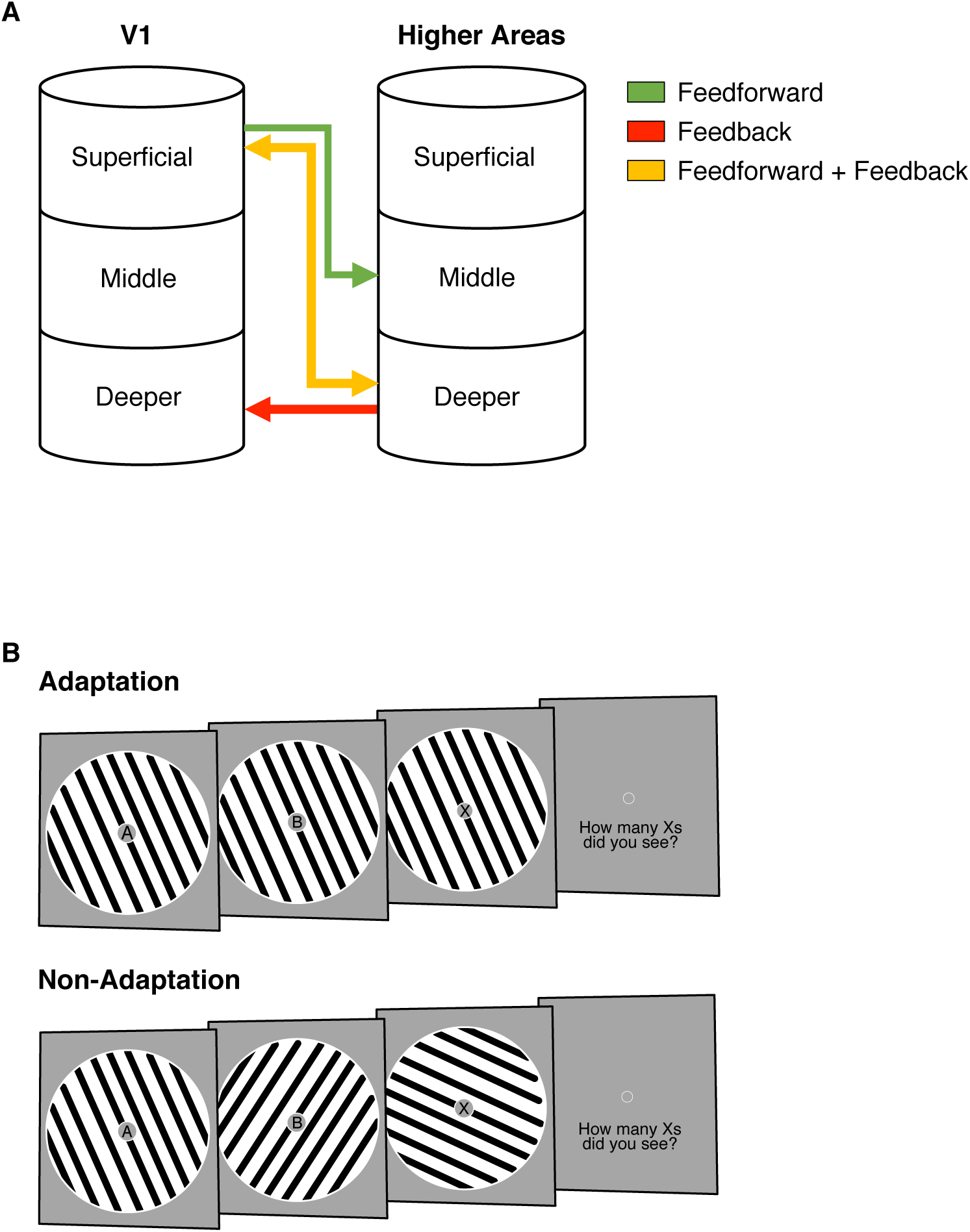
fMRI Laminar Circuits and fMRI Design. (A) Schematic representation of feedforward (superficial - middle layers; green), feedback (deeper - deeper layers; red), and feedforward plus feedback (superficial - deeper layers; yellow) anatomical connectivity between V1 and higher cortical regions. Here, we focussed on feedforward vs. feedback connections. (B) fMRI design. Adaptation blocks comprised 16 sinewave gratings presented at the same orientation. Non-adaptation blocks comprised 16 gratings presented at different orientations. During stimulus presentation, participants were asked to count the number of times a target letter (e.g., X) was displayed throughout the stream of distracters and report it at the end of each stimulus block.

## Results

### fMRI adaptation across cortical depth in visual cortex

To test whether adaptation alters visual orientation processing, we measured fMRI responses when participants (N=15) were presented with gratings (n=16 per block) either at the same orientation (adaptation) or different orientations (non-adaptation) in a blocked fMRI design (Figure 1B). Participants were asked to perform a Rapid Serial Visual Presentation task (RSVP; i.e. detect a target in a stream of letters presented in the centre of the screen) to ensure that they attended similarly across conditions (Larsson, Landy, & Heeger, 2006).

We tested for fMRI adaptation in visual cortex due to stimulus repetition by comparing fMRI responses for adaptation (i.e. the same oriented sinewave grating presented repeatedly within a block) vs. non-adaptation (i.e. gratings of varying orientation presented in a block). To test for differences in orientation-specific fMRI adaptation across cortical depth, for each participant we mapped the retinotopic areas in the visual cortex (V1, V2, V3, V4), assigned voxels in three layers (deeper, middle, superficial) using an equi-volume approach (*see Methods, MRI Data Analysis: Segmentation and Cortical Depth Sampling*; Figure 2D), and extracted fMRI responses across layers. To control for possible differences in thermal noise, physiological noise or signal gain across layers (Goense, Merkle, & Logothetis, 2012; Havlicek & Uludağ, 2020), we a) matched the number of voxels across layers for each participant and ROI and b) z-scored the laminar-specific time courses to control for differences in variance across layers, while preserving condition-dependent differences within each cortical layer.

**Figure 2.**
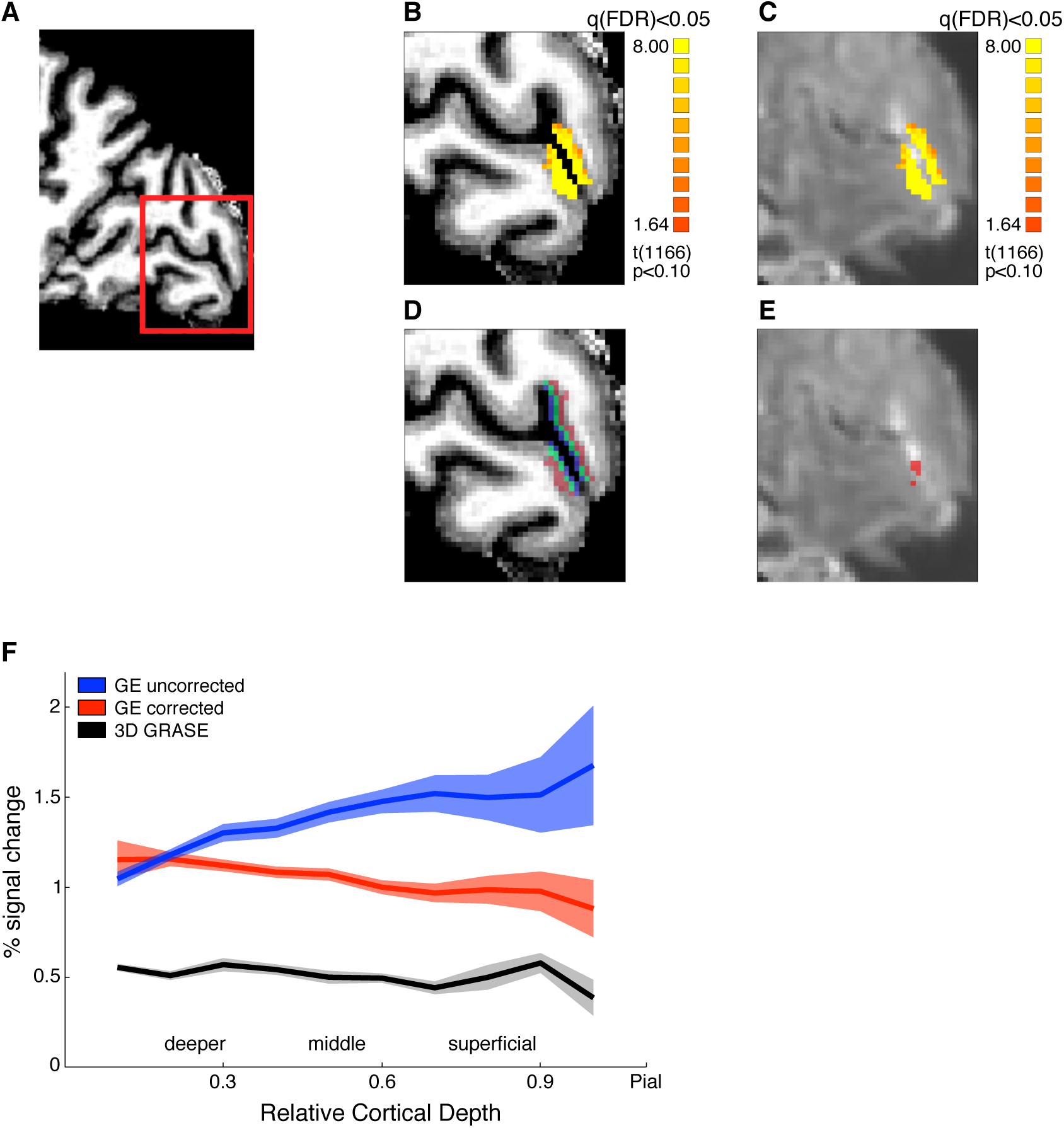
fMRI data analysis, layer segmentation, and vascular contribution correction. (A) Sagittal brain view of representative participant: red insert highlights region of interest (ROI, early visual cortex). Structural (B) and functional (C) images of the ROI showing activation maps for stimulus vs. fixation. Activation is well confined within the grey matter borders. (D) Mapping of cortical layers within the ROI: deeper layers shown in red, middle layers in green, and superficial layers in blue. (E) Voxels confounded by vasculature effects (in red) overlaid on mean functional image. (F) Mean BOLD (percent signal change from fixation baseline) across participants for V1 across cortical depth. Comparison between BOLD signal before (blue) and after tSNR and t-value correction (red), and 3D GRASE BOLD signal (black). The superficial bias observed in the BOLD signal is reduced after correction and matches closely the laminar profile of the 3D GRASE data.

Our results showed fMRI adaptation (i.e. decreased fMRI responses for adaptation compared to non-adaptation) across cortical layers in visual areas (Figure 3A; Figure 3 – supplement 1). In particular, a repeated measures ANOVA showed significant main effects of ROI (V1, V2, V3, V4; F(3,39)=23.276, *p*=0.001), layer (deeper, middle, superficial; F(2,26)=4.942, *p*=0.034), and condition (adaptation, non-adaptation; F(1,13)=11.872, *p*=0.004). There was no significant ROI x condition x layer interaction (F(6,78)=1.949, *p*=0.131), suggesting similar fMRI adaptation across visual areas. A significant condition x layer interaction (F(2,26)=9.506, *p*=0.002) indicated stronger fMRI adaptation in superficial than middle and deeper layers. Post-hoc comparisons showed significantly decreased fMRI responses for adaptation across cortical layers (deeper: t(14)=-3.244, *p*=0.006; middle: t(14)=-3.920, *p*=0.002; superficial: t(14)=-4.134, *p*=0.001).

**Figure 3.**
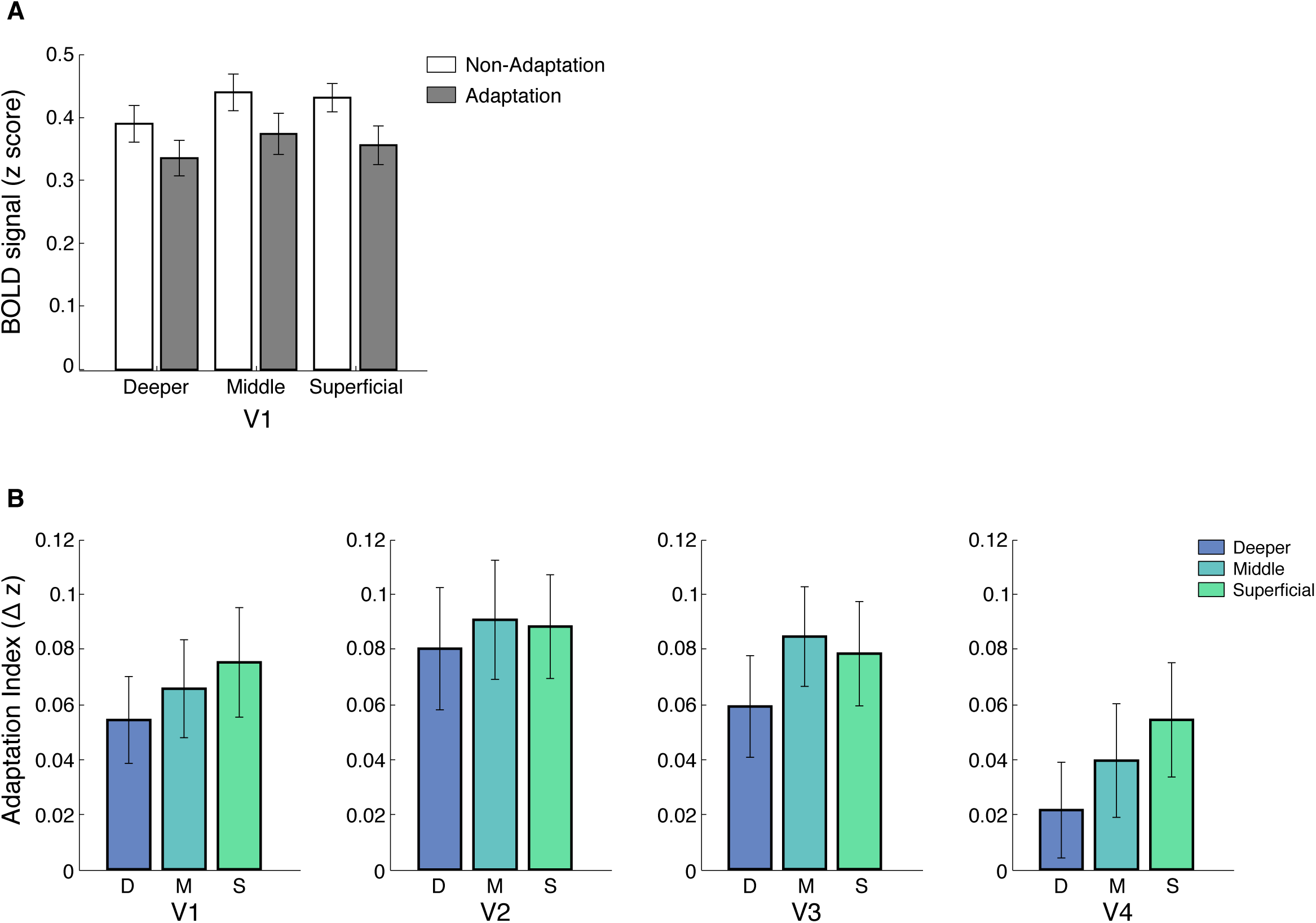
Laminar BOLD and Adaptation Index for V1, V2, V3, and V4. (A) Mean BOLD across V1 cortical layers. Bar-plot shows z-scored BOLD signal from fixation baseline for adaptation (grey) and non-adaptation (white) across cortical layers of V1. Error bars indicate standard error of the mean across participants (N=15). (B) fMRI adaptation index across cortical layers (D: deeper, M: middle, S: superficial) for V1, V2, V3, and V4. Bar plots show difference in z-scored BOLD signal between the non-adaptation and the adaptation conditions. Error bars indicate standard error of the mean across participants (N=15 for V1, V2, and V3; N=14 for V4).

We observed a similar pattern of results when we controlled for signal contribution from voxels at the border of adjacent layers, using a spatial regression analysis (Kok, Bains, Van Mourik, Norris, & De Lange, 2016; Koster, Chadwick, Chen, Hassabis, & Kumaran, 2018; Markuerkiaga, Barth, & Norris, 2016). To unmix the signal, we regressed out the time course of voxels assigned to middle layers and adjacent to the superficial layers from the time course of voxels assigned to superficial layers. We applied the same approach to voxels assigned to the deeper layers and adjacent to the middle layers. Figure 3A shows stronger fMRI adaptation in superficial V1 layers following this correction (see Figure 3 – supplement 1 for higher visual areas). We observed significantly decreased fMRI responses across layers (deeper: t(14)=-3.438, *p*=0.004, middle: (t(14)=-3.682, *p*=0.002, superficial: t(14)=-3.779, *p*=0.002) and a significant condition x layer interaction (F(2,28)=5.996, *p*=0.022).

To further quantify fMRI adaptation across cortical depth, we computed an fMRI adaptation index (i.e. fMRI responses for non-adaptation minus adaptation) per layer and ROI (Figure 3B). A repeated-measures ANOVA showed a significant main effect of layer (F(2,26)=9.506, *p*=0.002), ROI (F(3,39)=5.858, *p*=0.003) and no significant ROI x layer interaction (F(6,78)=1.949, *p*=0.131). Post-hoc comparisons showed significantly stronger fMRI adaptation in superficial compared to deeper (t(14)=-3.083, *p*=0.009) and middle compared to deeper cortical layers (t(14)=-3.821, *p*=0.002) in visual cortex.

### Control analyses

To control for potential confounds due to the contribution of vasculature-related signals to BOLD, we conducted the following additional analyses.

First, it is known that BOLD measured by GE-EPI is higher at the cortical surface due to vascular contributions (Ugurbil, Toth, & Kim, 2003; Uludag, Müller-Bierl, & Ugurbil, 2009; Yacoub, Van De Moortele, Shmuel, & Ugurbil, 2005). To ensure that the fMRI adaptation we observed in superficial layers was not confounded by this superficial bias, we identified and removed voxels with high temporal signal to noise ratio and high t-statistic for stimulation contrast (see Methods: *Regions of Interest (ROI) Analysis*). Figure 2F shows that following these corrections the superficial bias in the GE-EPI acquired BOLD was significantly reduced. That is, the magnitude and variance of GE-EPI BOLD signals from voxels closer to the pial surface were reduced, as indicted by a significant interaction between GE-EPI acquired BOLD signal from different cortical depths (deeper, middle, superficial) before vs. after correction (F(2,28)=58.556, *p*<0.0001). That is, the superficial bias corrections resulted in decreased BOLD signal mainly in middle and superficial layers as indicated by post-hoc comparisons (middle: t=7.992, *p*<0.0001; superficial: t=11.241, *p*<0.0001).

Second, we scanned a subset of participants (n=5) with a 3D GRASE sequence that is known to be sensitive to signals from small vessels and less affected by larger veins, resulting in higher spatial specificity of the measured BOLD signal (e.g., De Martino et al., 2013; Kemper et al., 2015). Consistent with previous studies (De Martino et al., 2013), the 3D GRASE data showed: a) overall lower BOLD signal in V1 compared to the GE-EPI acquired BOLD data and b) similar BOLD amplitude across V1 cortical depths. Figure 2F shows that the corrected GE-EPI BOLD signal in V1 follows a similar pattern across cortical depth as the 3D GRASE BOLD. In particular there were no significant differences in BOLD acquired with 3D GRASE vs. the corrected GE-EPI BOLD signal across cortical depths (i.e. no significant sequence x layer interaction: F(2,6)=2.878, *p*=0.187), suggesting that our superficial bias corrections reduced substantially the contribution of vasculature-related signals in GE-EPI measurements. Finally, we observed similar fMRI adaptation patterns across cortical layers for the data collected with 3D GRASE (Figure 3 – supplement 2), suggesting that the fMRI adaptation effects we observed in superficial layers could not be simply attributed to superficial bias effects.

Taken together our results demonstrate fMRI adaptation across layers in visual cortex with stronger effects in superficial than middle and deeper layers, suggesting stronger adaptation-related changes in read-out signals in superficial layers rather than input signals in middle layers. Further, comparing performance on the RSVP task during scanning across conditions showed that it is unlikely that these fMRI adaptation effects were due to differences in attention, as the RSVP task was similarly difficult across conditions. In particular, the mean performance across participants (adaptation condition: 62.7% ±0.3%; non-adaptation condition 60.15% ±0.4%, SEM) did not differ significantly between conditions (t(12)=0.312, *p*=0.76).

### fMRI Adaptation in intraparietal cortex

We next tested for adaptive processing in posterior parietal cortex regions (IPS1, IPS2 Benson et al., 2014, 2012; Wang et al., 2015) that have been shown to be involved in processing expectation due to stimulus familiarity (de Lange et al., 2018; Garrido et al., 2009; Li et al., 2010; Summerfield & De Lange, 2014). We observed fMRI adaptation across cortical layers in IPS1 and IPS2 (i.e. significantly decreased responses for adaptation than non-adaptation, Figure 4). In particular, a repeated measures ANOVA showed a main effect of condition (F(1,13)=7.994, *p*=0.013), layer (F(2,26)=25.824, *p*<0.0001), and no significant interactions between condition and layer (F(2,26)=0.575, *p*=0.511), nor between ROI, condition, and layer (F(2,26)=0.639, *p*=0.510). Spatial regression analysis showed similar pattern of results (main effect of condition: F(1,13)=6.149, *p*<0.05, cortical layer: F(2.26)=22.359, *p*<0.0001), suggesting similar fMRI adaptation effects across cortical layers in posterior parietal cortex.

**Figure 4.**
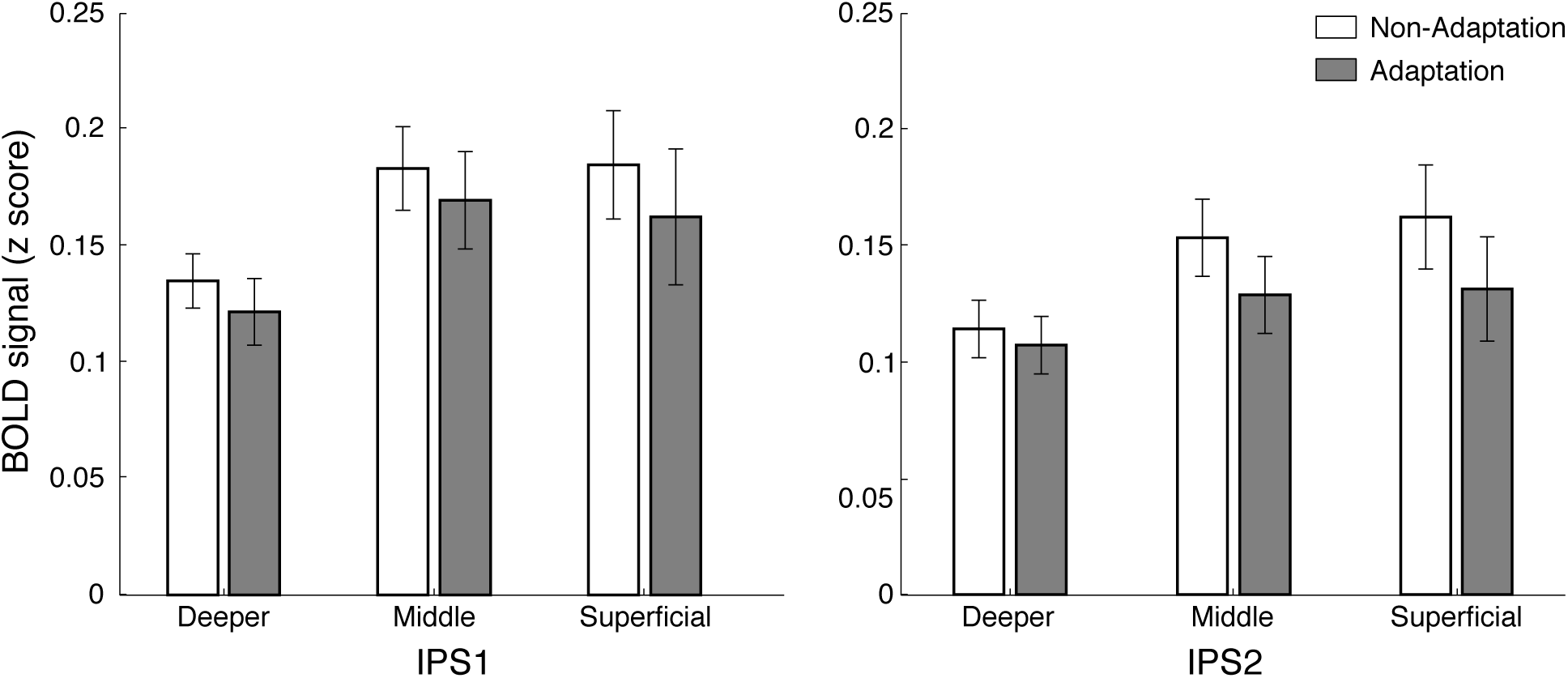
Laminar BOLD for IPS1, IPS2. Mean BOLD in IPS1 and IPS2 across cortical layers. Bar plots show z-scored BOLD signal for adaptation (grey) and non-adaptation (white) across cortical layers of IPS1 and IPS2. Error bars indicate standard error of the mean across participants (N=15).

### Functional Connectivity analysis

Ultra-high field fMRI allows us to interrogate the finer functional connectivity across areas based on known anatomical models of connectivity across cortical layers. Recent work (for review: Lawrence, Formisano, Muckli, & de Lange, 2019) has proposed that anatomical connections between superficial V1 layers and middle layers of higher areas relate to feedforward processing, while anatomical connections between deeper V1 layers and deeper layers of higher areas relate to feedback processing (Figure 1A). We tested functional connectivity in these circuits to discern feedforward vs. feedback processing for orientation-specific adaptation. We did not test connectivity between superficial V1 layers and deeper layers of higher areas, as these connections are known to relate to both feedback and feedforward processing.

We used ICA-based denoising and Finite Impulse Response functions (FIR) to denoise and deconvolve the time course data per layer, controlling for noise and potential task-timing confounds. Comparing functional connectivity (i.e. Pearson correlation between the eigenvariate time courses) within visual cortex and between visual and posterior parietal cortex showed dissociable results. That is, we observed stronger feedforward connectivity for adaptation within visual cortex (i.e. V1 superficial and V4 middle layers), while stronger feedback occipito-parietal connectivity (i.e. V1 deeper layers and IPS). A repeated measures ANOVA showed a significant three-way interaction (F(2,26)=5.089, *p*=0.031) between circuit (V1-V4, V1-IPS1, V1-IPS2), connectivity (feedforward, feedback), and condition (adaptation, non-adaptation).

In particular, for functional connectivity within visual cortex, we tested differences in connectivity between V1 and V4 layers that represent early and later stages of processing in the visual pathway. Our results showed significantly higher functional connectivity between V1 superficial layers and V4 middle layers for adaptation compared to non-adaptation (t(13)=3.03, *p=*0.01), suggesting enhanced feedforward processing for adaptation within visual cortex (Figure 5). In contrast, no significant differences between conditions were observed in functional connectivity between V1 deeper layers and V4 deeper layers, that is known to relate to feedback processing (t(13)=0.98, *p*=0.351).

**Figure 5.**
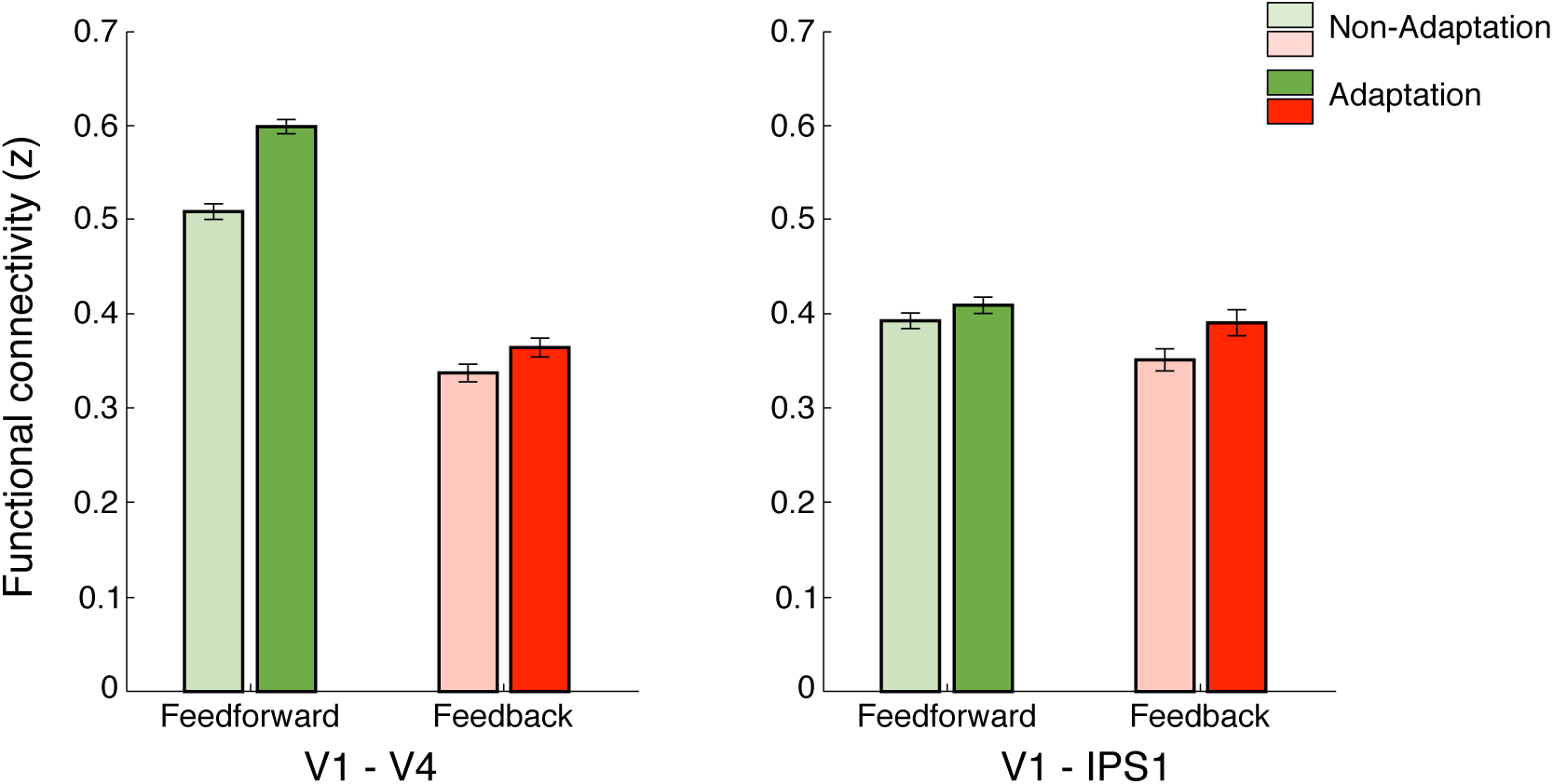
Functional Connectivity. Functional connectivity using Finite Impulse Response. Bar plots show Fisher z transformed r values for connectivity between V1 and V4 and V1 and IPS1. Feedforward anatomical connections are tested between: a) V1 superficial and V4 middle layers, b) V1 superficial layers and IPS1. Feedback anatomical connections are tested between: a) V1 and V4 deeper layers, b) V1 deeper layers and IPS1. Darker bars indicate adaptation, lighter bars indicate non-adaptation. Error bars indicate standard error of the mean across participants (N=14 for V1-V4; N=15 for V1-IPS1).

For occipito-parietal connectivity, we tested differences in connectivity between V1 layers and IPS subregions (IPS1, IPS2) as there were no significant differences in fMRI adaptation across IPS layers (F(2,28)=0.542, p=0.493). Our results showed significantly higher functional connectivity between V1 deeper layers and IPS1 for the adaptation compared to the non-adaptation condition (t(14)=2.15, *p*=0.049). This result is consistent with fMRI adaptation in V1 deeper layers (t(14)=3.438, *p*=0.004) and suggests enhanced feedback processing for visual adaptation. In contrast, no significant differences in functional connectivity between V1 superficial layers and IPS1 (i.e. functional connectivity related to feedforward processing) were observed between conditions (t(14)=0.7313, *p*=0.477). Further, no significant differences in functional connectivity between conditions were observed between V1 and IPS2 (V1 deeper layers and IPS2: t(14)=1.12, *p*=0.281; V1 superficial layers and IPS2: t(14)=0.551, *p*=0.59).

## Discussion

Here, we exploit UHF laminar fMRI to interrogate adaptive processing across cortical depth at a finer scale than afforded by standard fMRI methods. Our results advance our understanding of the brain mechanisms that mediate adaptive processing in the following main respects. First, visual adaptation is implemented by recurrent processing of signals in visual cortex, as indicated by fMRI adaptation (i.e. BOLD decrease due to stimulus repetition) across layers with stronger effects in superficial than middle and deeper layers. Second visual adaptation extends beyond the visual cortex, as indicated by decreased BOLD across layers in posterior parietal cortex. Third, dissociable connectivity mechanisms mediate adaptive processing: feedforward connectivity within the visual cortex, indicating inherited adaptation from early to higher visual areas (e.g., Larsson et al., 2016; Solomon & Kohn, 2014), while feedback connectivity from posterior parietal to visual cortex, reflecting top-down influences (i.e. expectation of repeated stimuli) on visual processing.

In particular, sensory inputs are known to enter the cortex at the level of the middle layer (middle layer 4) and output information is fed forward through the superficial layer (superficial layer 2/3). In contrast, feedback information is thought to be exchanged mainly between deeper layers (deep layer 5/6) (Larkum, 2013; Markov et al., 2014). Neurophysiological studies have shown that this micro-circuit is involved in a range of visual recognition (Self, van Kerkoerle, Supèr, & Roelfsema, 2013; Van Kerkoerle et al., 2014), and attention (Buffalo, Fries, Landman, Buschman, & Desimone, 2011) tasks. Recent laminar fMRI studies provide evidence for the involvement of this circuit in the context of sensory processing (De Martino et al., 2015) and visual attention (Fracasso, Petridou, & Dumoulin, 2016; Lawrence, Norris, & De Lange, 2019; Scheeringa, Koopmans, Van Mourik, Jensen, & Norris, 2016). Our results show fMRI adaptation across layers in primary visual cortex with stronger effects in superficial than middle and deeper layers, suggesting that adaptation alters more strongly the processing of read-out than input sensory signals. These signals are then forwarded to higher visual areas, as indicated by increased functional connectivity between V1 and V4. These results are consistent with previous neurophysiological studies showing that sensory adaptation is a fast form of plasticity (Gutnisky & Dragoi, 2008; Whitmire & Stanley, 2016; Xiang & Brown, 1998) and brain imaging studies showing that adapted BOLD responses in higher visual areas are inherited from downstream processing in V1 (Ashida, Kuriki, Murakami, Hisakata, & Kitaoka, 2012; Larsson et al., 2016).

It is important to note that despite the advances afforded by UHF imaging, GE-EPI remains limited by vasculature-related signals contributing to BOLD at the cortical surface, resulting in loss of spatial specificity (Kay et al., 2019). To reduce this superficial bias, we removed voxels with high temporal signal to noise ratio (Olman, Inati, & Heeger, 2007) and high t-statistic for stimulation contrast (Kashyap, Ivanov, Havlicek, Poser, & Uludağ, 2018; Polimeni, Fischl, Greve, & Wald, 2010). Further, we applied a signal unmixing method (Kok et al., 2016; Koster et al., 2018) to control for draining vein effects from deep to middle and middle to superficial layers. We compared BOLD signals across conditions (adaptation vs. non-adaptation) and layers after z-scoring the signals within each layer to account for possible differences in signal strength across layers (Goense, Merkle, & Logothetis, 2012; Havlicek & Uludağ, 2020). Following these corrections, we observed stronger fMRI adaptation (i.e. stronger BOLD response for non-adapted than adapted stimuli) in superficial layers, suggesting that our results are unlikely to be confounded by vasculature-related superficial bias. In addition, we corroborated our fMRI adaptation results in superficial layers using a 3D GRASE sequence that measures BOLD signals that are less affected by macro-vascular contribution. Our findings on orientation-specific adaptation in superficial layers are consistent with previous laminar imaging studies showing BOLD effects in superficial layers in a range of tasks (De Martino et al., 2015; Olman et al., 2012). Recent advances in cerebral blood volume (CBV) imaging using vascular space occupancy (VASO) (e.g., Beckett et al., 2019; Huber, Uludağ, & Möller, 2019) could be exploited in future studies to enhance the spatial specificity of laminar imaging in the human brain.

A possible mechanism by which orientation-specific adaptation is implemented in visual cortex is via recurrent processing of signals across V1 columns (Self et al., 2013). Horizontal connections across V1 columns are known to mediate iso-orientation inhibition (Malach, Amir, Harel, & Grinvald, 1993); that is suppression of neurons that are selective for the same orientation across columns. Iso-orientation inhibition is shown to be more pronounced in superficial layers and support orientation tuning (Rockland & Pandya, 1979). In particular, previous work has shown that horizontal connections between V1 columns primarily terminate in middle and superficial layers (Rockland & Pandya, 1979) and pyramidal cells in superficial layers make extensive arborisations within the same layer (Douglas & Martin, 2007). Consistent with this interpretation, previous neurophysiological studies have shown stronger decrease in neural population responses due to stimulus repetition in superficial layers of V1, while delayed adaptation effects in middle and deeper levels (Westerberg, Cox, Dougherty, & Maier, 2019). The neural connectivity within superficial V1 layers (layer 2/3) has been also shown to be rapidly and dynamically modulated by sensory adaptation (Hansen & Dragoi, 2011). In particular, before exposure to prolonged stimulus presentation, neurons tuned to the same orientation were shown to be strongly connected (i.e. cross-correlation between pairs of neurons). However, after adapting to a non-preferred orientation, stronger connectivity was observed between neurons tuned to the adapted orientation.

An alternative explanation is that BOLD effects in superficial layers reflect feedback processing (e.g. Gau, Bazin, Trampel, Turner, & Noppeney, 2020; Muckli et al., 2015). Previous work has shown that synaptic input to superficial layers may result due to increase in feedback signals carried by neurons that have dendrites projecting to the superficial layers and their cell bodies in deeper layers (Larkum, 2013). Our results showing fMRI adaptation in deeper layers in primary visual cortex and increased functional connectivity between IPS and deeper V1 layers suggest that long-range feedback from the posterior parietal cortex contributes to adaptive processing in V1, consistent with the role of parietal cortex in expectation and prediction due to stimulus repetition. Recent fMRI studies focusing on higher visual areas have investigated the role of expectation in repetition suppression that– similarly to sensory adaptation for simple stimulus features in V1– is characterized by decreased BOLD responses to more complex stimuli (i.e. faces, objects) in higher visual areas (Grill-Spector et al., 2006). In particular, Summerfield et al., (2008) showed stronger repetition suppression in the lateral occipito-temporal cortex for identical stimulus pairs that were repeated frequently, providing evidence for a role of top-down influences (i.e. expectation) in repetition suppression and visual processing.

A framework for linking adaptive processing within visual cortex and feedback repetition suppression mechanisms due to expectation is proposed by the predictive coding theory (Friston, 2005; Rao & Ballard, 1999; Shipp, 2016). According to this framework, perception results from comparing feedback expectation and prediction signals in upstream regions with feedforward signals in sensory areas. When these signals match, the error (i.e., the difference between the prediction fed back and the incoming sensory input) is low; in contrast, when the expectation does not match with the sensory input, the prediction error is high resulting in increased neural responses for unexpected compared to expected (i.e. repeated stimuli). Bastos et al., (2012) have proposed a microcircuit model of predictive coding that combines excitatory and inhibitory properties of pyramidal neurons across cortical layers to account for prediction encoding, prediction errors, and modulation of incoming sensory inputs to minimise prediction error. Considering our findings in light of this model provides insights in understanding the circuit underlying adaptive processing in the human brain. It is likely that long-range top-down information (e.g. expectation signals from posterior parietal cortex) is fed back to the deeper layers of V1 and it is then compared with information available at the superficial layers (i.e. iso-orientation inhibition). A mismatch (i.e. prediction error) of signals (i.e. expectation of a repeated stimulus compared to the presentation of an unexpected stimulus) results in decreased fMRI responses for expected compared to unexpected stimuli. This is consistent with previous laminar imaging studies showing stronger fMRI responses in deeper or superficial V1 layers for perceptual completion tasks and suggesting top-down influences in visual processing (Kok et al., 2016; Muckli et al., 2015).

In sum, exploiting UHF imaging, we provide evidence that adaptive processing in the human brain engages a circuit that integrates recurrent processing within visual cortex with top-down influences (i.e. stimulus expectation) from posterior parietal cortex via feedback. This circuit of local recurrent and feedback influences is critical for rapid brain plasticity that supports efficient sensory processing by suppressing familiar and expected information to facilitate resource allocation to new incoming input. Combining laminar imaging with electrophysiological recordings has the potential to shed more light in the dynamics of this circuit, consistent with recent evidence (Buffalo et al., 2011; Self et al., 2013; Van Kerkoerle et al., 2014) that gamma oscillations are linked to feedforward processing in input layers, while alpha/beta oscillations are related to feedback mechanisms in superficial and deeper cortical layers. Understanding these circuit dynamics is the next key challenge in deciphering the fast brain plasticity mechanisms that support adaptive processing in the human brain.

## Material and Methods

### Participants

Eighteen healthy volunteers (11 female, 7 male) participated in the study. Seventeen participants were scanned with a Gradient Echo-Echo Planar Imaging (GE-EPI) sequence (main experiment). Due to the lack of previous 7T fMRI adaptation studies, we determined sample size based on power calculations following a 3T fMRI study from our lab using the same paradigm (Karlaftis et al., 2019) that showed fMRI adaptation for effect size of Cohen’s f^2^=0.396 at 80% power. Data from two participants were excluded from further analysis due to excessive head movement (higher than 1.5mm) and technical problems during acquisition, resulting in data from fifteen participants for the main experiment (mean age: 24.44 years and SD: 3.83 years). Five participants (four who participated in the main experiment and an additional participant) were scanned with a 3D GRASE EPI sequence. All participants had normal or corrected-to-normal vision, gave written informed consent and received payment for their participation. The study was approved by the local Ethical Committee of the Faculty of Psychology and Neuroscience at Maastricht University.

### Stimuli

Stimuli comprised sinewave gratings (1 cycle/degree) of varying orientations (Figure 1B). Stimuli were presented centrally, within an annulus aperture (inner radius: 0.21 degrees; outer radius: 6 degrees). The outer edge of the aperture was smoothed using a sinusoidal function (standard deviation: 0.6 degrees). Experiments were controlled using MATLAB and the Psychophysics toolbox 3.0 (Brainard, 1997; Pelli, 1997). For the main fMRI experiment, stimuli were presented using a projector and a mirror setup (1920 × 1080 pixels resolution, 60 Hz frame rate) at a viewing distance of 99 cm. The viewing distance was reduced to 70 cm for the control experiment, as a different coil was used, and adjusted so that angular stimulus size was the same for both scanning sessions.

### Experimental Design

#### fMRI session

Both the main and control fMRI experiments comprised a maximum of 8 runs (13 participants completed 8 runs for each experiment; 2 participants in the main experiment and 1 participant in the control completed 6 runs). Each run lasted 5 min 6 s, and started with 14s fixation, followed by 6 stimulus blocks, three blocks per condition (adaptation, non-adaptation) and ended with 14s fixation. The order of the blocks was counterbalanced within and across runs. Each block comprised 16 stimuli followed by 2s for response to the RSVP task. The orientation of the gratings was drawn randomly from uniform distributions, ranging from -85° to -5°, and +5° to +85° in steps of 7.27°, excluding vertical (i.e., 0°). The same orientation was presented across adaptation blocks per participant. 16 different orientations were presented per block for the non-adaptation condition. Each stimulus was displayed for 1900ms with a 100ms inter-stimulus interval for both the adaptation and non-adaptation conditions to ensure similar stimulus presentation parameters (e.g. stimulus transients) between conditions. During scanning participants were asked to perform a Rapid Serial Visual Presentation (RSVP) task. A stream of letters was presented in rapid serial order (presentation frequency: 150ms, asynchronous to the timings of grating presentation) within an annulus at the centre of the screen (0.5 degrees of visual angle). Participants were asked to fixate at the annulus and report, by a key press at the end of each block, the number of targets (1-4 per block). No feedback was provided to the participants.

In the same scanning session, anatomical data and fMRI data for retinotopic mapping were collected following standard procedures (e.g., Engel, Glover, & Wandell, 1997).

### MRI acquisition

Imaging data were acquired on a 7T Magnetom scanner (Siemens Medical System, Erlangen, Germany) at the Scannexus Imaging Centre, Maastricht, The Netherlands. Anatomical data were acquired using an MP2RAGE sequence (TR = 5s, TE = 2.51ms, FOV = 208 x 208mm, 240 sagittal slices, 0.65 mm isotropic voxel resolution).

For the main experiment (n=17), we used a 32-channel phased-array head coil (NOVA Medical, Wilmington, MA, USA) and a 2D Gradient Echo, Echo Planar Imaging (GE-EPI) sequence (TE = 25ms, TR = 2s, voxel size = 0.8 mm isotropic, FOV = 148 x 148 mm, number of slices = 56, partial Fourier = 6/8, GRAPPA factor = 3, Multi-Band factor = 2, bandwidth = 1168 Hz/Pixel, echo spacing = 1ms, flip angle = 70°). The field of view covered occipito-temporal and posterior parietal areas; manual shimming was performed prior to the acquisition of the functional data.

For the control experiment (n=5), participants were scanned with a 3D inner-volume gradient and spin echo (GRASE) sequence with variable flip angles (Feinberg & Oshio, 1991; Kemper, De Martino, Yacoub, & Goebel, 2016). This sequence is largely based on a spin echo sequence for which the measured T2-weighted BOLD signal has higher spatial specificity and is less confounded by large draining veins near the pial surface (e.g., Duong et al., 2003; Goense, Zappe, & Logothetis, 2007; Kemper et al., 2015; Uludag, Müller-Bierl, & Ugurbil, 2009). We used a custom-built surface-array coil (Sengupta et al., 2016) for enhanced SNR of high-resolution imaging of visual cortex (TR = 2 s, TE = 35.41 ms, FOV = 128 x 24mm, number of slices = 12, echo-spacing = 1.01ms, total readout train time = 363.6, voxel size = 0.8 mm isotropic, 90° nominal excitation flip angle and variable refocussing flip angles ranging between 47° and 95°). The latter was used to exploit the slower decay of the stimulated echo pathway and hence to keep T2-decay-induced blurring in partition-encoding direction at a small, acceptable level, that is, comparable to the T2*-induced blurring in typical EPI acquisition protocols for functional imaging (Kemper et al., 2016).

### MRI data analysis

#### Segmentation and cortical depth sampling

T1-weighted anatomical data was used for coregistration and 3D cortex reconstruction. Grey and white matter segmentation was performed on the MP2RAGE images using FreeSurfer (http://surfer.nmr.mgh.harvard.edu/) and manually improved for the regions of interest (i.e., V1, V2, V3, V4, and IPS) using ITK-Snap (www.itksnap.org, Yushkevich et al., 2006). The refined segmentation was used to obtain a measurement of cortical thickness. Following previous studies, we assigned voxels in three layers (deeper, middle, superficial) using the equi-volume approach (Kemper, De Martino, Emmerling, Yacoub, & Goebel, 2018; Waehnert et al., 2014) as implemented in BrainVoyager (Brain Innovation, Maastricht, The Netherlands). This approach has been shown to reduce misclassification of voxels to layers, in particular for regions of interest presenting high curvature. Information from the cortical thickness map and gradient curvature was used to generate four grids at different cortical depths (ranging from 0, white matter, to 1, grey matter). Mapping of each voxel to a layer was obtained by computing the Euclidean distance of each grey matter voxel to the grids: the two closest grids represent the borders of the layer a voxel is assigned to (Figure 2D).

For the 3D GRASE control experiment, we used the LAYNII tools (https://github.com/layerfMRI/LAYNII), as they provided better segmentation for images with a limited field of view.

#### GE-EPI Functional data analysis

The GE-EPI functional data were analysed using BrainVoyager (version 20.6, Brain Innovation, Maastricht, The Netherlands) and custom MATLAB (The MATHWORKS Inc., Natick, MA, USA) code. Preprocessing of the functional data involved three serial steps starting with correction of distortions due to non-zero off-resonance field; that is, at the beginning of each functional run, five volumes with inverted phase encoding direction were acquired and used to estimate a voxel displacement map that was subsequently applied to the functional data using COPE, BrainVoyager, Brain Innovation. The undistorted data underwent slice-timing correction, head motion correction (the single band image of each run was used as reference for the alignment), high-pass temporal filtering (using a GLM with Fourier basis set at 2 cycles) and removal of linear trends. Preprocessed functional data were coaligned to the anatomical data using a boundary-based registration approach, as implemented in BrainVoyager (Brain Innovation, Maastricht, The Netherlands). Results were manually inspected and further adjusted where needed. To validate the alignment of functional to anatomical data, we calculated the mean EPI image of each functional run for each region of interest (ROI) and estimated the spatial correlation between these images (e.g., Marquardt, Schneider, Gulban, Ivanov, & Uludağ, 2018). We performed manual adjustment of the alignment if the spatial correlation was below 0.85. We excluded a small number of runs (n=3, n=1 for two participants respectively), as their alignment could not be improved manually.

#### 3D GRASE functional data analysis

Functional images were analysed using BrainVoyager (version 21.0, Brain Innovation, Maastricht, The Netherlands), custom MATLAB (The MATHWORKS Inc., Natick, MA, USA) code, and Advanced Normalization Tools (ANTs, Avants et al., 2011) for images registration. The first volume of each run was removed to allow for the magnetisation to reach a steady state. Head motion correction was performed using as reference the first image (10 volumes with TR=6s) acquired at the beginning of the functional runs. The higher contrast of this image facilitated the coregistration of the anatomical and functional images. After motion correction, temporal high-pass filtering was applied, using a GLM with Fourier basis set at 3 cycles per run. Preprocessed images were converted into Nifti files and an initial manual registration was performed between the first image and the anatomical image using the manual registration tool provided in ITK-Snap (www.itksnap.org, Yushkevich et al., 2006). The resulting transformation matrix was applied to coregister the anatomical image to the functional space and fine-tuned adjustments were provided by means of antsRegistration tools.

#### Regions of Interest (ROI) analysis

We used the data from the retinotopic mapping scan to identify regions of interest. For each participant, we defined areas V1 to V4 based on standard phase-encoding methods. Participants viewed rotating wedges that created travelling waves of neural activity (e.g., Engel et al., 1997). Due to limited coverage during acquisition, area V4 was identified for 14 of the 15 participants. Intraparietal regions (comprising IPS1, and IPS2) were defined for each participant based on anatomical templates provided by Benson (https://hub.docker.com/r/nben/occipital_atlas/; Benson, Butt, Brainard, & Aguirre, 2014; Benson et al., 2012; Wang, Mruczek, Arcaro, & Kastner, 2015). This procedure uses the individual participant-based segmentation obtained with Freesurfer and an anatomical probabilistic template, to estimate the best location for the region of interest (i.e. IPS). Each IPS subregion was subsequently inspected to ensure consistent definition across participants. For each ROI and individual participant, we modelled BOLD signals using a GLM with two regressors, one per stimulus condition (adaptation, non-adaptation). We included estimated head motion parameters as nuisance regressors. The resulting t-statistical map was thresholded (t=1.64, *p*=0.10) to select voxels within each ROI that responded more strongly to the stimulus conditions compared to fixation baseline (Figure 2B, C).

Voxel selection within each ROI was further refined by excluding voxels that were confounded by vasculature effects that are known to contribute to a superficial bias in the measured BOLD signal; that is, increased BOLD with increasing distance from white matter (see Results: *Control Analyses*). In particular, it has been shown that the BOLD signal measured using GE-EPI, T2* weighted sequences is confounded by vasculature-related signals (Ugurbil et al., 2003; Uludag et al., 2009; Yacoub et al., 2005) due to veins penetrating the grey matter and running through its thickness, as well as large pial veins situated along the surface of the grey matter (Duvernoy, Delon, & Vannson, 1981). This results in increased sensitivity (i.e., strong BOLD effect) but decreased spatial specificity of the measured signal.

Here, we took the following approach to reduce superficial bias due to vasculature contributions. First, following previous work (Olman et al., 2007), we computed the temporal signal to noise ratio (tSNR) for each voxel in each ROI (V1, V2, V3, V4, IPS). We used this signal to identify voxels near large veins that are expected to have large variance and low intensity signal (mean tSNR across V1 smaller than 12.02 ± 2.02) due to the local concentration of deoxygenated haemoglobin resulting in a short T2* decay time. Second, it has been shown that high t-values on a fMRI statistical map are likely to arise from large pial veins (Kashyap et al., 2018; Polimeni et al., 2010). Therefore, voxels with t-score values above the 90^th^ percentile (mean t-score across V1 larger than t=15.47 ± 4.16.) of the t-score distribution obtained by the GLM described above, were removed from further analysis.

Further to account for possible differences in signal strength across cortical layers due to thermal and physiological noise, as well as signal gain (Goense, Merkle, & Logothetis, 2012; Havlicek & Uludağ, 2020), we a) matched the number of voxels across layers (i.e. to the layer with the lowest number of voxels) per participant and ROI, and b) z-scored the time courses within cortical layer per ROI, controlling for differences in signal levels across layers while preserving signal differences across conditions (after correction of vascular contributions, e.g. Lawrence, Norris, & De Lange, 2019). Normalised fMRI responses for each condition (adaptation, non-adaptation) were averaged across the stimulus presentation (excluding participant responses; 32 - 34s after stimulus onset), blocks, and runs for each condition. For visual cortex ROIs we focused on the time window that captured the peak of the haemodynamic response to visual stimulus presentation (4 to 18s after stimulus onset). We conducted repeated measures ANOVAs to test for significant differences between conditions (adaptation, non-adaptation), cortical depth (deeper, middle, superficial layers) and ROIs (V1, V2, V3, V4, IPS1, IPS2). Pairwise t-test comparisons were used as post-hoc tests.

#### Functional Connectivity analysis

We followed standard analyses methods to compute functional connectivity across ROIs and layers. We pre-processed the functional and anatomical data in SPM12.3 (v6906; http://www.fil.ion.ucl.ac.uk/spm/software/spm12/). We first performed brain extraction and normalisation to MNI space on the anatomical images (non-linear). The functional images were then corrected for distortions, slice-scan timing (i.e. to remove time shifts in slice acquisition), head motion (i.e. aligned each run to its single band reference image), coregistered all EPI runs to the first run (rigid body), coregistered the first EPI run to the anatomical image (rigid body) and normalised to MNI space (applying the deformation field of the anatomical images). Data were only resliced after MNI normalisation to minimize the number of interpolation steps.

Next, we used an ICA-based denoising procedure (Griffanti et al., 2014). We applied spatial smoothing (2mm) and linear detrending, followed by spatial group ICA. The latter was performed using the Group ICA fMRI Toolbox (GIFT v3.0b) (http://mialab.mrn.org/software/gift/). Principal Component Analysis (PCA) was applied for dimensionality reduction, first at the subject level, then at the group level. A fixed number (n=35) of independent components was selected for the ICA estimation. The ICA estimation (Infomax) was run 20 times and the component stability was estimated using ICASSO (Himberg, Hyvärinen, & Esposito, 2004). Group Information Guided ICA (GIG-ICA) back-reconstruction was used to reconstruct subject-specific components from the group ICA components (Du et al., 2016). The results were visually inspected to identify noise components according to published procedures (Griffanti et al., 2017). We labelled 12 of the 35 components as noise that captured signal from veins, arteries, CSF pulsation, susceptibility and multi-band artefacts.

To clean the fMRI signals from signals related to motion and the noise components, we followed the soft cleanup approach (Griffanti et al., 2014) on the BrainVoyager unsmoothed data in native space (see *GE-EPI Functional data analysis*). That is, we first regressed out the motion parameters (translation, rotation and their squares and derivatives; Friston, Holmes, Poline, Price, & Frith, 1996) from each voxel and ICA component time course. Second, we estimated the contribution of every ICA component to each voxel’s time course (multiple regression). Finally, we subtracted the unique contribution of the noise components from each voxel’s time course to avoid removing any shared signal between neuronal and noise components.

Further, following recent work (Cole et al., 2019), we deconvolved the denoised time courses using Finite Impulse Response functions (FIR). In particular, we fitted 23 regressors per condition that covered the duration of each task block, including the response period and fixation block, to capture the whole hemodynamic response. This method allows us to accurately model and remove the cross-block mean response for each condition (adaptation, non-adaptation) to account for potential task-timing confounds that have been shown to inflate the strength of the computed task-based functional connectivity. Within the GLM, the data were high-pass filtered at 0.01Hz and treated for serial autocorrelations using the FAST autoregressive model (Corbin, Todd, Friston, & Callaghan, 2018; Olszowy, Aston, Rua, & Williams, 2019). For each ROI and layer, we then computed the first eigenvariate across all voxels within the region to derive a single representative time course per layer and ROI for connectivity analysis. We computed functional connectivity as the Pearson correlation between the eigenvariate time courses across ROIs and layers. Finally, we performed a paired t-test on the functional connectivity values (after Fisher z-transform) to test for significant differences in connectivity between conditions (adaptation, non-adaptation).

## Acknowledgements

We would like to thank Christopher Wiggins and Esther Steijvers (Scannexus) for technical support, Peter Kok (University College London), Denis Schluppeck (University of Nottingham), Federico De Martino (University of Maastricht), Laurentius Huber (University of Maastricht), and Cheryl Olman (University of Minnesota) for the expert and insightful comments to the manuscript. We would also like to thank Adrian Ng, Valentyna Chernova, and Cher Zhou for help with the analysis. This work was supported by grants to Z.K. from the Biotechnology and Biological Sciences Research Council (H012508 and BB/P021255/1) and the Wellcome Trust (205067/Z/16/Z).

---

**Figure 3 - Supplement 1:**
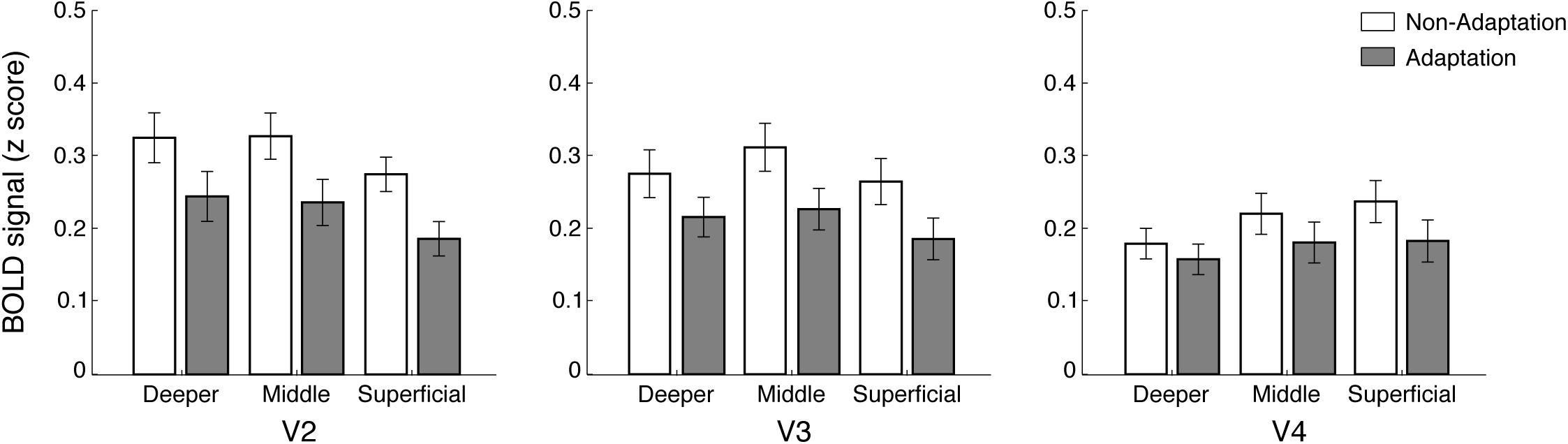
Laminar BOLD in higher visual areas. Mean laminar BOLD in V2, V3 and V4 cortical layers. Bar-plots show z-scored BOLD signal change from fixation baseline for adaptation (grey) and non-adaptation (white) across cortical layers in V2, V3, and V4. Error bars indicate standard error of the mean across participants (N=15 for V2 and V3; N=14 for V4).

**Figure 3 - Supplement 2:**
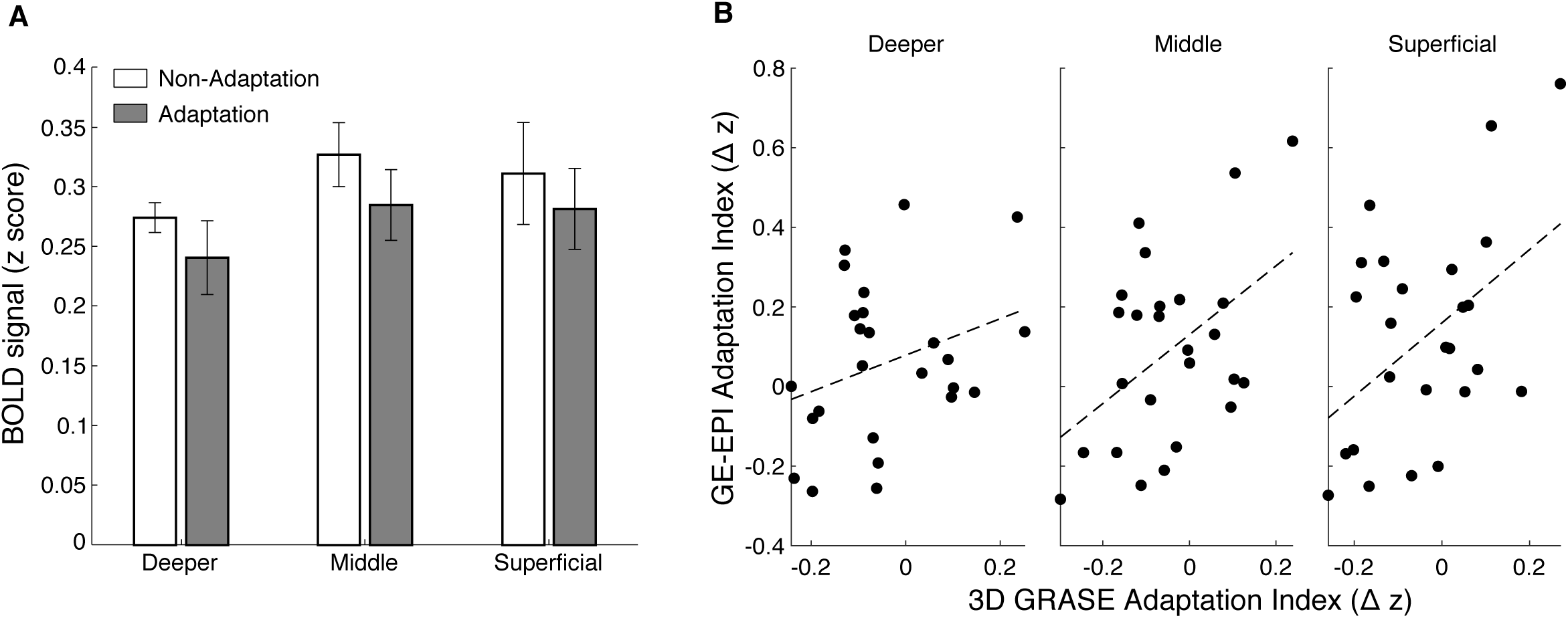
3D GRASE fMRI. (A) Mean z-scored BOLD (signal change from fixation baseline) measured with 3D GRASE for the adaptation (grey) and non-adaptation (white) across cortical layers in V1. Decreased BOLD for the adaptation, compared to the non-adaptation condition can is shown across cortical layers. The difference between conditions was not statistically significant (condition x cortical layer: F(2,6)=0.875, *p*=0.432) due to the small sample size (N=4). (B) Correlation of fMRI adaptation index for participants scanned with both GE-EPI and 3D GRASE sequences (N=4) across cortical layers. Individual dots indicate fMRI adaptation index (z-scored BOLD for non-adaptation minus z-scored BOLD for adaptation), for each participant and run (N_runs_=5 for 1 participant; N_runs_=6 for 1 participant; N_runs_=8 for 2 participants) for GE-EPI (y-axis) and 3D GRASE (x-axis) sequences. Black dashed lines indicate correlation between GE-EPI and 3D GRASE fMRI adaptation index. Correlations were stronger in superficial (r=0.465, *p*=0.019) and middle (r=0.467, *p*=0.019), than deeper (r=0.315, *p*=0.125) layers. This correspondence of results across sequences suggests that our results showing stronger fMRI adaptation in superficial than middle and deeper layers, could not be simply due to vasculature confounds.

